# The contribution of the ankyrin repeat domain of TRPV1 as a thermal module

**DOI:** 10.1101/641803

**Authors:** E. Ladrón-de-Guevara, G. E. Rangel-Yescas, D. A. Fernández-Velasco, A. Torres-Larios, T. Rosenbaum, L. D. Islas

## Abstract

The TRPV1 cation non-selective ion channel plays an essential role in thermosensation and perception of other noxious stimuli. TRPV1 can be activated by low pH, high temperature or naturally occurring pungent molecules such as allicin, capsaicin or resiniferatoxin. Its noxious thermal sensitivity makes it an important participant as a thermal sensor in mammals. However, details of the mechanism of channel activation by increases in temperature remain unclear. Here we used a combination of approaches to try to understand the role of the ankyrin repeat domain (ARD) in channel behavior. First, a computational modeling approach by coarse-grained molecular dynamics simulation of the whole TRPV1 embedded in a phosphatidylcholine (POPC) and phosphatidylethanolamine (POPE) membrane provides insight into the dynamics of this channel domain. Global analysis of the structural ensemble shows that the ankyrin repeat domain is a region that sustains high fluctuations during dynamics at different temperatures. We then performed biochemical and thermal stability studies of the purified ARD by means of circular dichroism and tryptophan fluorescence and demonstrate that this region undergoes structural changes at similar temperatures that lead to TRPV1 activation. Our data suggest that the ARD is a dynamic module and that it may participate in controlling the temperature sensitivity of TRPV1.

**Statement of Significance:** This work demonstrates that the temperature-dependent dynamics of the ankyrin repeat domain (ARD) of TRPV1 channels, as probed by coarse-grained molecular dynamics, corresponds to the experimentally determined dynamics of an isolated ARD domain. These results show that this region of TRPV1 channels undergoes significant conformational change as a function of increased temperature and suggest that it participates in the temperature-dependent structural changes that lead to channel opening.

## Introduction

TRPV1 is a non-specific cation channel implicated in nociception by chemicals, temperature, and pH (1–3). This channel is one of the chemosensors involved in the sensation of pain and thermal stimuli and it participates in a diverse range of cellular processes (2, 4). The latter has been evidenced from studies where deletion of TRPV1 in mice alters noxious and mild temperature sensation (5, 6), while knock-out of other thermoTRPs such as TRPV2, TRPV3, and TRPV4 shows little effects in sensory transduction in rodents (6–8). Moreover, while deletion of TRPV1 in rodents does not affect corporal temperature, blockage of TRPV1 in vivo triggers hyperthermia (9).

The rat TRPV1 structure is a tetramer (Fig. 1a), with every monomer consisting of 838 amino acids (Fig. 1c). The structure solved by cryo-EM is from a minimal-functional TRPV1 that lacks 100 amino acids from the N-termini and 80 amino acids in the C-termini and which is also missing a longer S5-pore extracellular loop named the turret. This minimal-functional 586 amino acid construct provides a model for the full-length channel, although without unfolded loops. In TRPV1, the structure of the membrane-embedded domains is canonical with other ion channels like voltage-gated potassium, sodium and calcium ion channels. A tetramer is formed by a voltage sensing domain (VSD)-like domain, surrounding a pore formed by the contribution of the four pore domains (PD) of each subunit.

**Figure 1.**
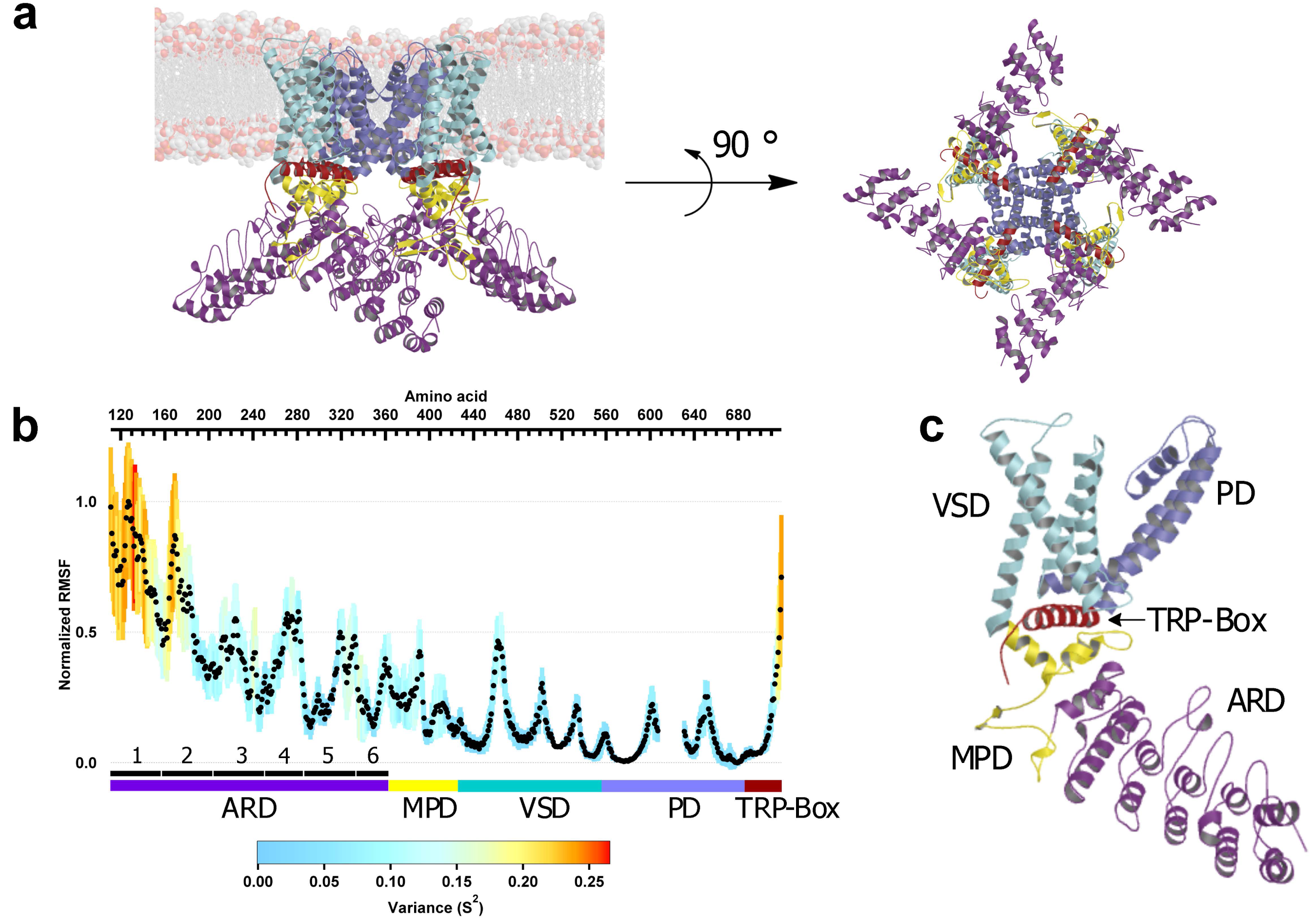
Depiction of the all-atom model of TRPV1, derived from the 3J5P PDB structure. (a, left), side view of TRPV1. The coloring of its domains is as follow: ARD, violet; MPD, yellow; VSD, cyan; PD, blue TRP-box, red. The bilayer is represented as spheres. (a, right), the intracellular view of the channel is represented with the same coloring code. Inter-subunit interactions are seen between MPD and ARD. (b) Analysis of the structural fluctuations induced by temperature by RMSF of the beads that correspond to main chain of each amino acid in coarse-grained MD. Averaged values were plotted as black dots. The discontinuity in the PD corresponds to the turret loop, which is not resolved in the cryo-EM structure. The ankyrin motifs have the greatest variance and RMSF are ANK1 and ANK2. All the ankyrin motifs are numbered from 1-6. (c) Single subunit depiction, voltage sensor-like domain (VSD) and pore domain (PD) are membrane embedded domains, ARD is fully cytosolic. The membrane proximal domain (MPD) has complex interactions with ARD, VSD, TRP-box. The TRP-box is sandwiched between the MPD and VSD.

The C-terminus of TRPV1 has been shown to be functionally important. Several modulating lipids, such as PIP2 and LPA bind in this region (10, 11). In the cryo-EM structures, the alpha helix after the S6 transmembrane segment is well defined and is known as the TRP-box; it is a conserved domain in almost all the members of the TRP family. In all the solved structures for TRPV1 (12–14), TRPV 2 (15), TRPV3 (16), TRPV4 (17), TRPV5 (18) and TRPV6 (19), the TRP-box is folded in the same manner. In the case of TRPV1, Cryo-EM experiments show that the TRP-box displays a conformational change when the channel is bound to capsaicin (13), or resiniferatoxin and double-knot toxin (DkTx) together (13), similar to what happens in TRPV2 sensitized by 2-Aminoethoxydiphenyl Borate (2-APB) (15). Until now the structure of the distal part of the C-terminus remains unknown, but there is some evidence that it is located near the ankyrin repeat domain (ARD), and that these two structures may undergo intradomain interactions (20).

The ARD and the membrane proximal domain (MPD), a structure located between the ARD and the first transmembrane segment, have a non-canonical interaction only seen in TRP channels. A “finger” loop from ANK4 touches the MPD at a recognition site in a neighboring subunit (Figure 1a and Sup. Fig. 1) (12, 19). Although some of the functions of these domains are still not known, others have been well characterized and include the direct interaction of the ARD with calmodulin (18, 21) and the possible participation of the MPD in heat activation (22). Thus, the ARD and MPD are functional regulatory domains and it should be noted that, before the first cryo-EM structures for TRP channels were obtained, the intracellular domains were visualized as individual and not interconnected domains. Thanks to structural analysis, today we know that the intracellular domains of TRPV ion channels, form tetramers and that coupling between these domains regulates channel function (19, 23). In this context, the overall nature of ARD interactions is poorly understood, but some evidences correlate with the notion that they play a leading role in the functional mechanism of the channel (21, 24). For example, there is some evidence of the ARD contributing to gating by selective oxidation of intracellular cysteines present in the ARD (3, 25). Additionally, a complex formed by TRPV5 and calmodulin was solved Recently (18). This interaction can be of interest since it also exists in TRPV1 and other members of the family and it is located between C-termini and the ARD.

To further study the role of the ARD in TRPV1 we studied the integral relationships between ARD function and channel gating. For this purpose, we employed coarse-grained molecular dynamics of the whole TRPV1 channel embedded in a POPC/POPE membrane and also performed experiments with the purified ARD in solution, making use of the thermal shift unfolding, circular dichroism, and tryptophan fluorescence measurements. We find that the ARD is a dynamical domain and that these dynamics are modulated by temperature in a way that might be relevant to activation by temperature.

## Methods

### Model Preparation

The cryo-electron microscopy (cryo-EM) structure of a minimal-functional rat TRPV1, solved at 3.4 Å resolution, was used as initial coordinates to perform coarse-grained (CG) molecular dynamics (PDBID: 3J5P) (12). The missing loops and side chains were built in the Swiss Modeler web server (26, 27). To generate our model, the server used the whole channel model (12) and the transmembrane cryo-EM structure solved in nanodiscs (14). The repaired TRPV1 structure was prepared for coarse-graining by means of the Martinize.py (28) program. A 20 nm per side, square POPC/POPE (coarse-grained) membrane was prepared in CHARMM-GUI (29), the channel model was embedded into the membrane using Pymol scripts written in-house. The TRPV1-POPC/POPE structure was prepared for molecular dynamics (MD) simulation in the Gromacs 4.5 Suite (30, 31). The model was minimized and solvated with CG water and surrounded by 30 mM NaCl ions. A 200 ns NVT equilibration dynamics with a Berendsen Thermostat was performed before the production run (28, 30).

### Structural comparison

Composite alignment was performed for cryo-EM models of open and closed states of TRPV1 and TRPV3 ion channels using the TopMatch Server (32). A blue to red gradient was used to represent root-mean square deviation (RMSD) divergence in structural alignments. The Pymol module colorbyrmsd.py was used for coloring.

### Molecular Dynamics Simulations

We used coarse-grained molecular dynamics (CGMD) simulations to characterize the dynamics of the ion channel in the membrane. We simulated the initially repaired APO model (PDBID: 3J5P) at various temperatures varying from 290 K to 350 K (17 °C to 77 °C) sampling every 5 K. The structure of TRPV1 was embedded in a POPC/POPE bilayer with a MARTINI force field; several equilibrating simulations with temperature coupling were performed, and then an equilibrating simulation with pressure and temperature was performed until the RMSD was stable at 298 K (28). A production run at 290 K was sampled as an initial structure. From a stable point in a 290 K run, thirteen replicas were prepared. The RMSD corresponds to the mean of the whole protein main-chain over the whole simulation time and the root-mean square fluctuation (RMSF) corresponds to the mean of a single amino acid position over time. Both were calculated with Gromacs 4.5. The output trajectories were used as input for Normal Mode Analysis (NMA). This analysis was performed in bio3D for the four subunits (33). Production runs were performed in the “Miztli” HP Cluster Platform 3000SL supercomputer at the Supercomputing Facility of the National Autonomous University of México (UNAM).

### ARD expression and purification

The DNA sequence encoding for the ARD of rat TRPV1 (residues 100-262) was cloned into a pET28c vector (Merck-Millipore, USA). For expression, the Escherichia coli BL21 (DE3) strain (NewEngland Biolabs, USA) was transformed with this vector. Batch cultures of transformed cells were grown in Luria Bertani broth (Sigma-Aldrich, USA) to an OD of 0.7-0.8, protein expression was then induced with Isopropyl-β-D-1-thiogalactopyranoside 0.1 mM (GoldBio, Canada) for 12 h at room temperature. The cell pellet was lysed in Tris 50 mM, pH 8, dithiothreitol 0.1 mM, NaCl 300 mM (Sigma-Aldrich, USA). The lysate was purified by Histrap FF columns (GE Healthcare) with an imidazole gradient from 0 to 0.5 M. The eluted protein was concentrated and desalted in a PD-10 column (GE Healthcare) against a Tris 50 mM, pH 8, dithiothreitol 0.1 mM buffer. Then, the protein was injected into a cation exclusion chromatography column (MonoQ) and eluted against Tris 50 mM, pH 8, dithiothreitol 0.1 Mm buffer with an increased linear gradient from 0-1 M of NaCl. The final step was a size exclusion chromatography in a hand-made 275 mL Superdex 75 Column (Dimensions 2.50 × 60 cm, GE Healthcare). ARD concentration was determined using a theoretical extinction coefficient of 1.21 L/mg·cm obtained using the ProtParam Expasy WebServer (34) and by bicinchoninic acid assays with reduction compatible reagents (Pierce Chemicals, USA).

### Thermal Shift Unfolding

Assays were prepared in a 96 well plate for real-time PCR; each well was prepared with 1 μL of SYPRO Dye 1X (Sigma-Aldrich, USA), 1 μL of 1mg/mL solution of pure ARD, and 1 μL of solubility and stability II kit (Hampton Research, USA), water was added to fill 10 μL per well. A real-time thermocycler QuantStudio 3 (ThermoFisher, USA) was used for collecting data from 20 to 80 °C, sampling every 1 °C. The data were exported and analyzed in IgorPro 6 (WaveMetrics, USA).

### Size Exclusion Chromatography - Multiple Angle Light Scattering (SEC-MALS)

The SEC-MALS analysis was performed on a DAWN HELEOS multi-angle light scattering detector, with eighteen angles detectors and a 658.9 nm laser beam, (Wyatt Technology, Santa Barbara, CA, USA) and an Optilab T-rEX refractometer (Wyatt Technology) in-line with a Superdex 75 10/300 GL (GE Life Sciences) size exclusion chromatography analytical column. Experiments were performed using an isocratic pump (Agilent) with a flow of 0.5 ml/min at room temperature (25°C). Data collection was performed with ASTRA 6.1 software (Wyatt Technology). For the experiments, 300 µl at 1 mg/ml protein were loaded on the columns with running buffer of 20 mM glycine pH 9.5, 200 mM NaCl. The molecular weight (MW) and the ratio of gyration (Rg) were calculated by the ASTRA software.

### Circular Dichroism Spectroscopy

Circular dichroism (CD) spectroscopy was carried out in a JASCO J-715 spectropolarimeter with a 1 mm path length cuvette, the sample was prepared at 250 μg/mL in 5 mM sodium phosphate buffer. Samples were heated at a 0.5 K min^−1^ rate with a Peltier device (JASCO). A thermal unfolding curve was generated by measuring a spectrum for every step. CD data were analyzed in custom-made scripts and plotted with Igor Pro 6 (WaveMetrics, USA). Normalized ellipticity data obtained at 208 nm were fitted to the following sigmoid function to calculate the T_m_:

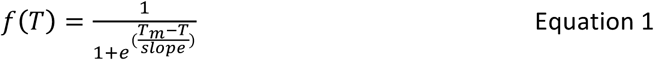

### Tryptophan Fluorescence Spectroscopy

Intrinsic fluorescence spectra were obtained in a PC1 spectrofluorometer (ISS Inc., USA) configured with a 10 W halogen lamp as the excitation source. All samples were prepared at 50 μg/L. Tryptophan fluorescence spectra were obtained using an excitation wavelength of 295 nm, emission spectra were collected from 310 to 400 nm, using excitation and emission slits of 0.25 and 1 mm respectively. The temperature was controlled with a Peltier device (Quantum Northwest). Normalized intensity at 330 nm and SCM data (see below) were fitted to Eq. 1.

The fluorescence spectral center of mass (SCM) was obtained from each spectrum and plotted as a function of temperature. The SCM was calculated from intensity data (I_λ_) obtained at different wavelengths (λ).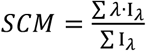

## Results

### Cytosolic domains of TRPV1 fluctuate more at higher temperature

It is generally assumed that TRPV1 is formed by a cluster of functional domains (Fig 1c), each with a set of specific interactions. The ARD, MPD and TRP-Box domains are intracellular. Several groups have found that the minimal functional TRPV1 structure needs the N-termini from the ANK1 motif upwards (12, 20, 35) and no more than one hundred amino acids from the C-terminus can be deleted without rendering the channel non-functional (35).

As an initial point in our research to understand the behavior of TRPV1 as a function of temperature, we carried out coarse-grain molecular dynamics simulations at different temperatures using a full-length model of TRPV1 embedded in a phosphatidylcholine/phosphatidylethanolamine (POPC/POPE) lipid membrane (Fig. 1a). The initial model was derived from the closed TRPV1 structure (PDBID: 3J5P) solved by cryo-EM (12). The model was simulated for 200 ns in temperature-controlled conditions (NVT) for stabilization and then 200 ns of pressure and temperature controlled conditions (NPT) with Berendsen thermostat and pressure coupling (see Methods). The simulations were equilibrated until the channel was in a stable RMSD state, from that point on, 12 replicas were simulated at 290, 295, 300, 305, 310, 315, 320, 325, 330, 335, 340, 345 and 350 K. At these temperatures independent runs of 500 ns were simulated to sample for an adequate time in order to observe critical large-scale structural transitions.

In figure 1b, we represent the RMSF of a single subunit of the whole TRPV1 tetramer. Temperatures from 290 to 350 K were sampled and averaged; each point is the average of RMSF of a particular residue for all the temperatures. The variance (s^2^) of the mean RMSF is plotted as a measure of dispersion from 0 to 0.26. The mean RMSF for all the subunit is 0.27 ± 0.22; 0.45 ± 0.21 for the ARD and 0.40 ± 0.21 for the ARD + MPD. The RMSF deviation, which correlates the fluctuation per amino acid during the trajectory of the dynamics, shows that there are some regions that fluctuate more than others as the temperature is increased. The regions with well-defined secondary structure have less movement and the loops and less structured chains move more (Figure 1b). The transmembrane region of the channel has smaller magnitude movements in relation to the cytosolic part; this is likely due to the two-dimensional restriction of the lipid membrane. Interestingly, the data shows that the mean RMSF greatly increases in the N and C-termini, even though these are regions with well-defined secondary structure.

All the N-terminus (ARD + MPD) shows a significant increase in main movement as temperature rises. The ankyrin motifs (ANK) are represented in Fig 1a. ANK1 and ANK2 are not solved in the cryo-EM structure (Supplemental Fig. 1c) and in our analysis these repeats presented a large average RMSF of 0.66 and the highest variance per amino acid, above 0.20. While the C-terminus also has a high value of temperature-dependent RMSF, for the purpose of this study, it will not be discussed.

### Protein dynamics and inter-subunit interactions of TRPV1

In order to study the coarse-grained dynamics of TRPV1 and provide a quantitative interpretation of the simulations, we performed normal mode analysis (NMA), (36, 37). We identified a first mode in which the ARDs undergo alternate up-and-down movements. Figure 2a and b shows the structure at two different stages of the oscillation. The arrows indicate the direction of the movement of each of the coarse-grained beads. The bottom panels show a view from the intracellular region of the channel. This normal mode represents more than 70% of the movements of each trajectory and this is seen at all the simulated temperatures.

**Figure 2.**
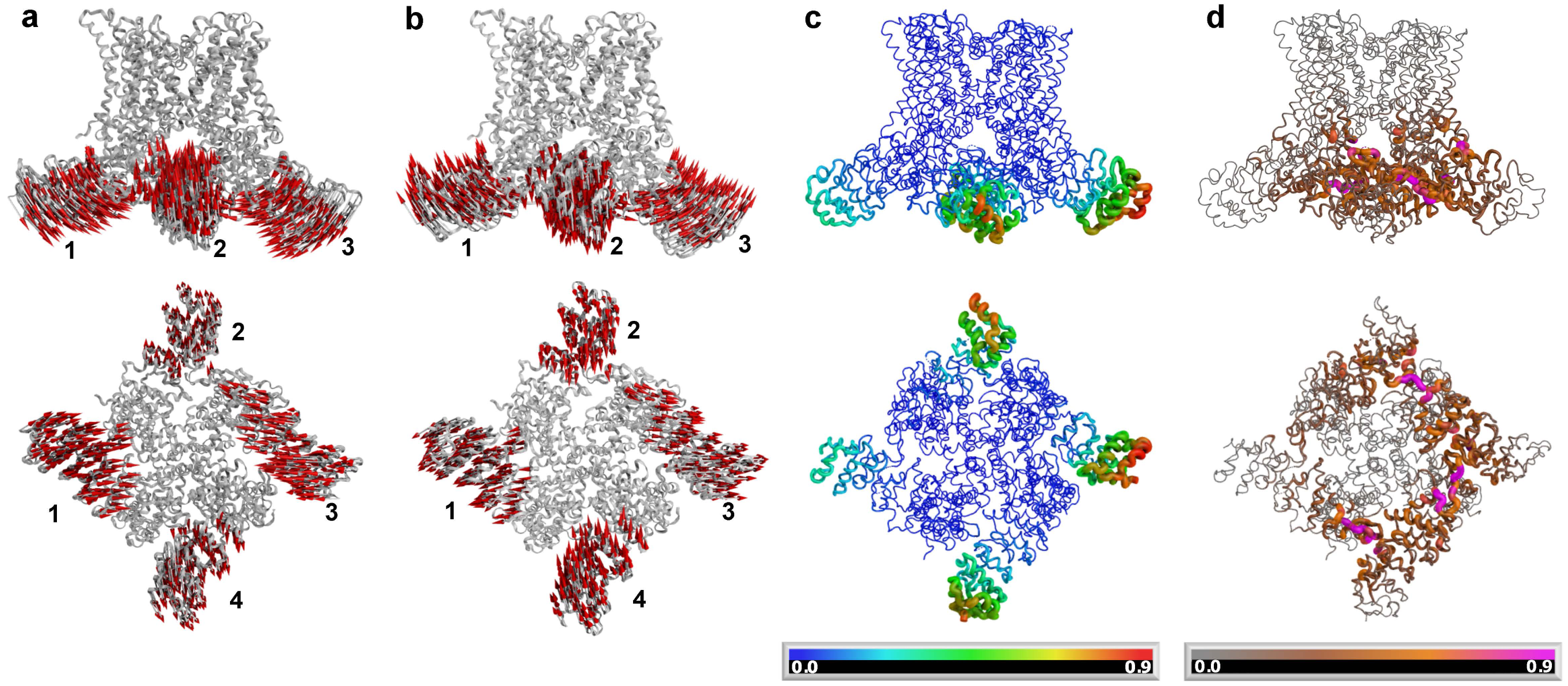
Normal mode analysis of TRPV1. The 290 K trajectory was analyzed to obtain normal modes. (a and b) The first mode shows an alternating upwards and downwards movement of adjacent ARDs. The red arrows indicate the direction of movement. The subunits are numbered for clarity. (a) Domains 1 and 3 moved downwards while 2 and 4 are upwards. (b) Opposed domains moved upwards (1 and 3) and downwards (2 and 4). The bottom panels show the intracellular view of the channel after a 90 ° rotation. (c) Fluctuations calculated by RMSF per bead relative to main chain. ARDs show the highest values of fluctuations. Bottom panel, view of a 90° rotation of the channel. The scale represents the value of the calculated fluctuation. (d) Deformation values of motion relative to neighbors. The MPDs show the highest correlated movement with the ARD (pink color). The values were calculated with Bio3d (see methods), (33, 36). The scale represents the calculated deformation values. The proximal ARD (ANK3 to MPD) was highly correlated.

Fluctuation analyses were performed from the trajectories by means of the greater NMA eigenvectors (33). Figure 2c shows the fluctuation analysis of the 290 K trajectory depicted from the same normal mode from 2a and 2b. It can be seen that the highest values of the fluctuations occur at the ankyrin repeats one and two and the fluctuations extend into the whole ARD.

A deformation displacement analysis was performed from the NMA; this is obtained by the averaged values of each individual mode, and by measuring the contribution of each backbone bead to the energy density as a function of position (36). This analysis provides information on the correlation between regions with high deformation and regions that are most affected by this deformation. Figure 2d shows that the regions with higher deformation are the ARD and that the regions with highest correlation with the deformation of the ARD is the MPD. This is a reflection of the physical interactions between these two domains but also means that the conformational changes in one domain are transmitted to the other in the interaction pair.

### Comparision of Molecular Dynamics Trajectory with CryoEM Structures

The structures of closed and open TRPV1 were compared by means of composite alignment. In contrast to rigid body alignment, this method allows the part-by-part alignment leaving out the regions without structural match.

The structure of TRPV1 in the apo, closed-like state (PDBID: 2J5P), the capsaicin-bound (CB) structure (PDBID: 2J5R) and an open-like structure bound to the activating double-knot toxin (DkTx) and resiniferatoxin (RTX) (PDBID: 2J5Q), were analyzed. First, the published DkTx/RTX toxin-bound structure (TXB) has a selectivity filter in an expanded open conformation (13) while the CB structure only shows an increase in the diameter of the lower gate (12, 13) (Supplemental figure 2). The closed-like conformation has similarity with both TXB and CB structures; in supplemental figure 2 we show that the closed structure is similar to CB in the upper part of transmembrane domain, as well as in the selectivity filter. The TXB structure has a wide-open selectivity filter but this structure is similar in the lower part of the transmemebrane domain, S6 helix and TRP-box domain to the CB structure. TRPV3 closed (PDBID: 6DVW) and open (PDBID: 6DVY) structures were also analyzed for comparison (Supplemental Fig 2c). The composite alignment of open vs. closed TRPV3 is closer to CB-TRPV1 vs. closed TRPV1 (23), (Supplemental Fig 2a and 2c). Closed TRPV1 and open TRPV3 are not comparable, but the composite alignment between structures has many similarities and importantly the ARD domain presents a high RMSD.

### The conformation of the ARD shows a thermal dependency at physiological temperature

The simulation data illustrates that the intracellular domains of TRPV1 undergo conformational changes at temperatures that are relevant to the physiological activation of the channel. The region that shows the largest mobility is the ARD, a 260 amino acid region in the N-terminus of TRPV channels. In TRPV1, the ARD is implicated in adenosine triphosphate binding, Ca^2+^-calmodulin modulation and regulation by cysteine oxidation (3, 21, 25, 38). In the whole channel, the ARD has several complex interactions with the pre-S1 region (MPD, 90 amino acids) in the same subunit, and a characteristic ARD-MPD interaction with the adjacent subunit (Fig 1a and 1c) (12). The ARD also has an interaction with the distal C-terminus, this region is not well solved in TRPV1 cryo-EM structure but in TRPV2 and TRPV3 it is (15, 16). The functional significance of these interactions remains unknown.

As a first approximation to study the biophysical properties of the N-terminus, we purified an ARD from rTRPV1 and screened its stability by a thermal shift assay (Figure 3), utilizing several pH and salt concentration conditions (see Methods). Figure 3a shows the fluorescence signal from the dye SYPRO orange as a reporter of the thermal unfolding process. SYPRO emits a fluorescence signal by binding to hydrophobic regions exposed during unfolding. The fluorescence signals show a characteristic sigmoid dependence on temperature, which is a feature of a two-state unfolding process in soluble proteins. To characterize the cooperativity of the change in fluorescence, the first derivative was calculated for every trace (Fig. 3a, 3b and 3c).

**Figure 3.**
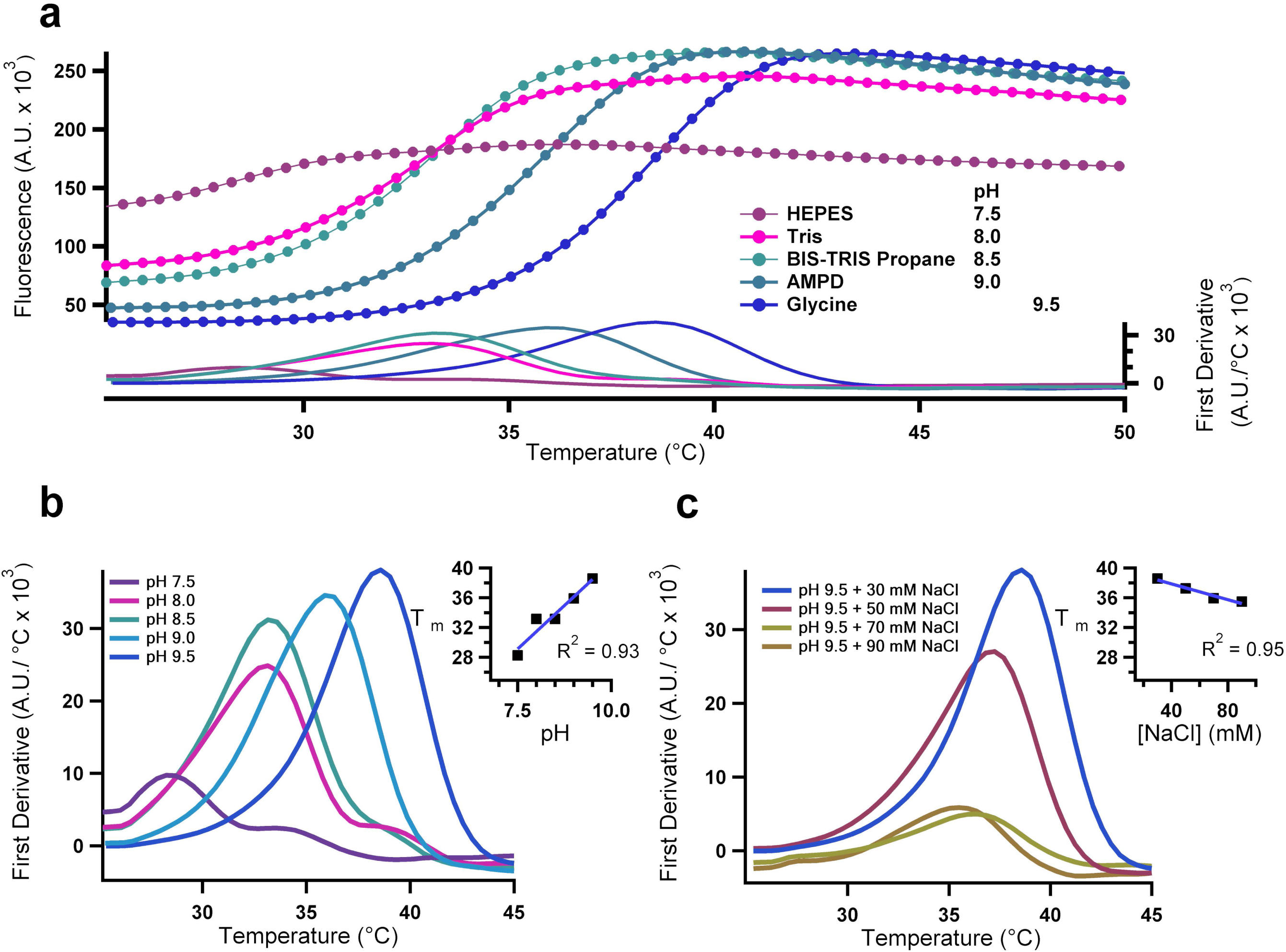
Thermal shift unfolding of ARD. (a) Representative raw SYPRO fluorescence signals obtained in a real time thermocycler are plotted as a function of temperature. The lower panel shows the first derivative of each signal. (b) First derivative (FD) plot (A.U./°C) of similar data as in (a) obtained at different pH values. The melting temperature, T_m_, was calculated as the temperature where the maximum occurs. Inset shows the T_m_ of the ARD at different pH values. (c) Melting temperature T_m_ of ARD in glycine buffer (pH 9.5) at different sodium chloride concentration. The inset shows the dependence of T_m_ on the NaCl concentration.

The data show that the first derivative of the fluorescence increases linearly as a function of pH. The data in Figure 3a, inset, indicate that the cooperativity of the unfolding of the ARD is highest at basic pH, as the value of the derivative increases as pH increases. The inset in Fig. 3b shows that additionally to the increased cooperativity, the domain becomes more stable, since the T_m_ value, defined as the temperature at which the derivative is maximal, shifts to higher temperatures. Next, the dependence on salt concentration was analyzed for the pH 9.5 condition. In Figure 3c we show the data for glycine buffer (pH 9.5) and several NaCl concentrations. The steepness of the fluorescence change is higher at low salt concentrations and the Tm decreases with increasing salt concentration, with a slope of −0.05 and a change in T_m_ from 36 to 38 °C. The behavior of ARD in glycine buffer (pH 9.5) was also studied by SEC-MALS to control for its possible oligomeric assembly. The experiment shows that the ARD is monodisperse in these conditions (Supplementary figure 3).

Significantly, these series of experiments indicate that the thermal transitions of the isolated ARD occur in a range of temperatures that is rather narrow (T_m_ = 36-38 °C for the most stable condition) and very close to the physiological temperature of mammals.

### Stability of the ARD in solution

Since SYPRO thermal shifts are mostly qualitative, we compared the thermal sensitivity of the ARD by means of tryptophan fluorescence and circular dichroism (CD) experiments. Figure 4a shows far-UV CD spectra (200 to 240 nm) of ARD obtained at increasing temperatures (see methods for details). The CD spectrum at 25°C is consistent with the high alpha helical content of the ARD structure. The shape and ellipticity of individual spectra changes as temperature is increased, indicating the presence of thermal unfolding of the ARD.

**Figure 4.**
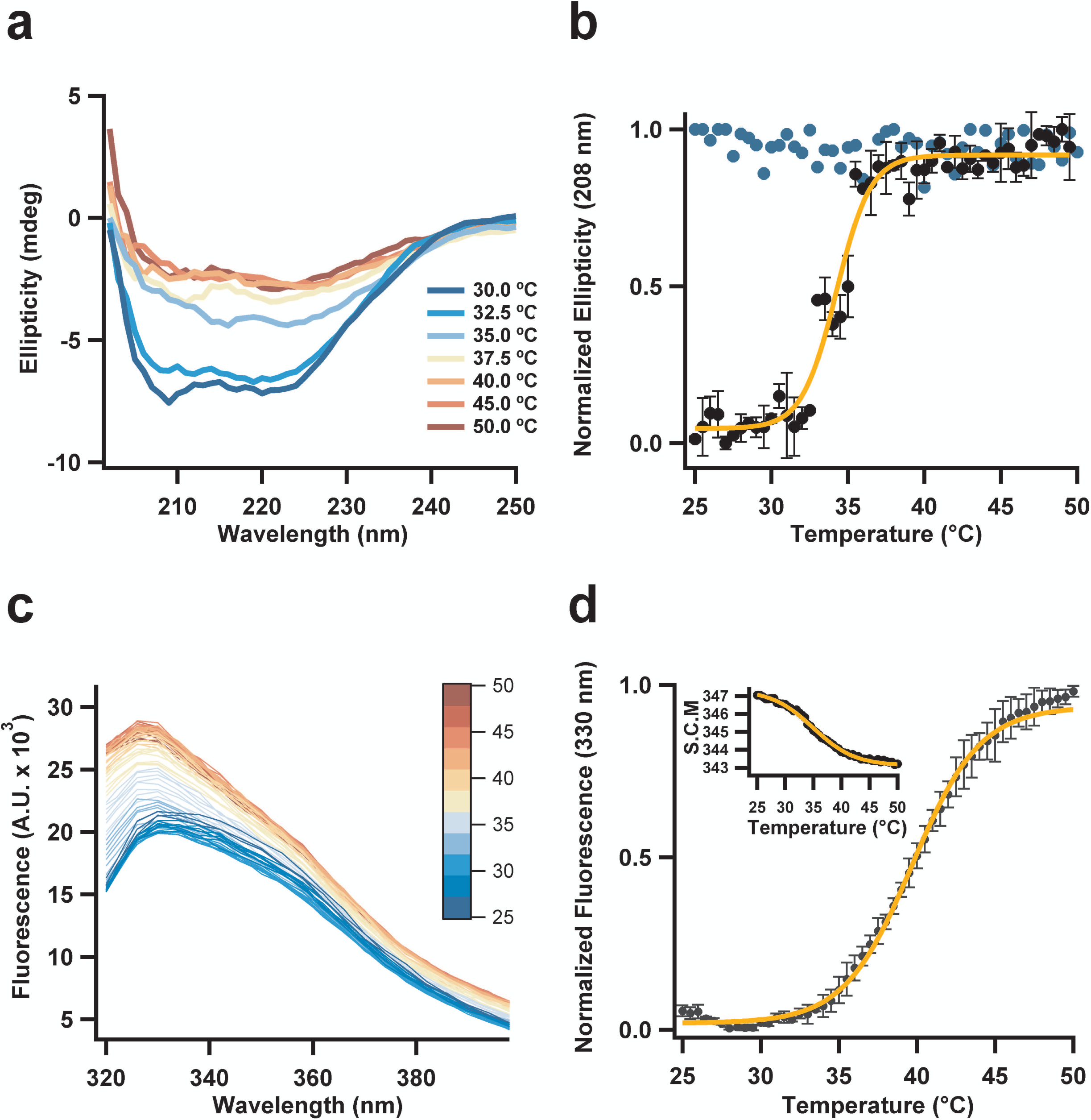
Thermal stability of ARD. (a) CD spectra of the ARD at the temperatures indicated in the inset f. (b) The normalized ellipticity change at 208 nm is plotted as a function of temperature. Data are mean and s.e.m. for n = 3 experiments (black circles). The blue circles are the data from one experiment at an attempt at refolding of the ARD by returning the temperature to 30 °C and indicate an irreversible unfolding transition. The continuous curve represents a fit to a sigmoid function (slope 1.13 ± 0.14). The value of T_m_ from the fit is of 34.2 °C ± 0.2 °C. (c) Tryptophan fluorescence spectra of the ARD at the temperatures indicated by the color scale. (d) The normalized fluorescence intensity increase for n = 3 experiments is plotted as the mean ± s.e.m. The continuous curve represents a fit to a sigmoid function (slope 2.17 ± 0.10). The value of T_m_ from the fit of intensity data is 39.7 °C ± 0.1 °C. The SCM analysis is shown in the inset. The parameters of the SCM fit are: slope 3.69 ± 0.13 and a T_m_ 35.0 ± 0.2 °C.

Since the main CD signal is due to a macro dipole formed by the sum of dipoles of all the secondary structures, the decrease of ellipticity is characteristic of a random array of secondary structures, even when a single alpha helix does not have cero signal. In the case of the ARD, the loss of signal can be associated to an increase of mobility between motifs. To characterize the change in CD, we normalized the signal at 208 nm and plotted it as a function of temperature (Fig. 4b). The unfolding occurs with a very steep dependence on temperature, with a T_m_ of ~34 °C, which is in agreement with the T_m_ observed in the thermal shift experiments. We attempted to measure the reversibility of the unfolding reaction; this result is plotted as blue circles in figure 4b and shows that thermal unfolding of the ARD is irreversible. In comparison, the closest ARD previously studied is from the hTRPV4, which has a T_m_ of 37.1 °C measured by CD but its reversibility was not tested (38).

Thermal unfolding was also measured by tryptophan fluorescence experiments. The ARD has an exposed Trp at position 272 of the ANK4 repeat. Excitation of this Trp at 295 nm and measurement of the fluorescence at 330 nm as a function of increasing temperature shows an increase in the fluorescence counts (Fig. 4c). The fluorescence increases (dequenching) in a temperature dependent manner and a sigmoid function was fitted with a T_m_ of 39.7°C (Fig 4d). A more robust analysis of Trp fluorescence dequenching is provided by a spectral center of mass analysis (SCM), which takes into account the contribution of the whole emission spectrum (39). This analysis is shown in the inset in figure 4d and indicates that the Tm value obtained from the fit is T_m_ of 34.9°C, which is closer to the T_m_ obtained from the CD experiments.

## Discussion

A defining characteristic of TRPV and other thermoTRP channels is the presence of several ankyrin repeats in the N-terminus. Elucidation of their functional role has been a main objective of TRP channel structure function studies. Others ARDs from different proteins show a variety of functions, from replication/translation regulation to protein-protein interaction and ligand binding (40–42).

TRPV1 is one of the best-studied members of the thermoTRP channel subfamily. Notwithstanding the wealth of available information, the mechanisms of heat sensitivity and heat-dependent activation remain poorly understood. Several mechanisms of thermal sensing have been proposed: The transmembrane regions could contribute with a significant heat capacity and serve as a thermal sensor domains (43); the pore domain could act as a temperature sensing domain (44) and the intracellular and transmembrane domains could act cooperatively to contribute to thermal sensation through a mechanism involving a linker domain (22). In TRPA1 channels, the ARD, although much bigger than the TRPV1 ARDs, have been implicated in temperature sensation.

Several TRPV1 structures determined by cryo-EM exist and are useful in interpreting experiments and proposing new ones, but the quality of these structures is in general poor. All of them are missing electronic density that should correspond to the initial ankyrin repeats of the ARD. Possible reasons for this are: high density of side chain of the amino acids, presence of salt bridges or the loss of density near Asp and Glu residues (45). Another possibility is higher mobility of this whole region of the protein structure (Supplemental figure 1). Our coarse-grain MD simulations with a repaired model show that indeed the amino terminus and, in particular the ARD, are highly dynamic regions. An important result stemming from these simulations is that increased temperatures also modulate the high mobility of ARDs. Experimentally, deletion of the ARD results in non-functional TRPV1 channels (12, 35), a result that has hindered an understanding of the role of ARs in heat sensitivity and other forms of gating in this channel. The ARDs are responsible for auto recognition, and probably an auto modulation mechanism by self-interaction.

Our approach identifies the dynamics of the ARD of TRPV1 as important for the overall function of the ion channel. The low frequency movement represented by the first normal mode identified in this study, suggest that the ARD is a dynamical module that might contribute to the response of TRPV1 to temperature. We propose that these fluctuations might be important in controlling temperature dependent gating of TRPV1. At the very least, our results highlight the importance of the ARD and its interactions with other parts of the channel in regulating channel gating.

The structure of the ARD of TRPV1 was previously solved by crystallography (46). Lishko et al. also showed that basic pH is better for ARD purification (46). Also, there are some groups that report a brief activation of the functional TRPV1 at basic pH (47).

Our data from CGMD simulations support a highly dynamical conformation of the ARD, and show breaking of some contacts between the second and third AR motifs at higher temperatures. This kind of perturbations may decrease stability of ARD as illustrated by the unfolding experiments in designed ARDs (40, 48) and might contribute to a conformational change coupled to channel opening.

Our biochemical data provide experimental support for the fluctuations observed in the simulations. Importantly, we observe conformational changes in the structure of the isolated ARD that occur at physiologically relevant temperatures. Our results indicate that the structure of the ARD undergoes important changes over a range of ~25°C. The CD data indicates that at 50 °C the structure is highly altered, but still contains a significant amount of alpha-helical structure. This result might indicate that the ARD is not completely unfolded into a random coil. An important observation is the fact that this structure is not reversible. In accordance with this, we have recently shown that temperature-dependent gating of full length TRPV1 is also irreversible and this irreversibility is reflected in a large hysteresis during the activation processes (49). It is thus possible that the behavior of the ARD observed here could be related to this irreversible gating process.

Some naturally occurring splicing variants of TRPV1 have been described. The ARD and MPD are part of two different exons, and the splicing variants ARD-less and MPD-less express ion channels that are not sensitive to temperature (50). In our simulations and structural analysis, it is observed that the distal ARD (ANK1-2) approaches the MPD domain. This interaction is hard to measure by MD because it is known that in this region a C terminal peptide forms a third beta strand of the MPD finger loop. Our analysis of the correlations of fluctuation during a simulated trajectory also highlights this interaction. Our data derived from NMA indicate that the regions that show highest correlated movement are precisely the proximal ARD and the MPD.

The importance of the ARD-MPD interaction in TRPV1 function is also highlighted by experiments by Yao et al., in which several TRPV1-MPD chimaeras were generated between rTRPV2, hTRPV2, mTRPV3 and mTRPV4. All the chimeras increased the sensitivity to temperature. These results are consistent with the ARD being in direct contact with the adjacent MPD, in such a way that its ability to interact with a viable MPD might play a role in the capacity of the TRPV channels to being thermally sensitive.

In conclusion, structure-function studies and natural occurring variants of the N-terminus (ARD + MPD) support its fundamental role for TRPV1 function. Our study contributes to an understanding of the dynamics of ARD and suggests that it participates in the regulation of thermal sensing, although its function as a thermal sensor or an element in temperature coupling remains to be elucidated.

## Author’s contributions

E. L-de-G, Designed experiments, performed experiments and simulations, analyzed data and wrote paper. G. E. R-Y, contributed materials, performed molecular biology and read the paper. D. A. F-V, helped with CD and fluorescence experiments, read the paper and analyzed data. A. T-L, helped with SECS-MALS and thermal shift experiments, read the paper and analyzed data. T. R., designed experiments and wrote the paper. L. D. Islas, Designed experiments, analyzed data and wrote the paper.

## Acknowledgments

E. Ladrón-de-Guevara is gratefull to Consejo Nacional de Ciencia y Tecnología (CONACYT) for a PhD scholarship (grant no. 297659). We acknowledge support from DGAPA-PAPIIT grant No. IN209515 to L.D.I and No. IN200717 to T.R. We also acknowledge support from CONACyT-Ciencia Básica grant No. 252644 to L.D.I and No. A1-S-8760 to T.R. We are specially gratefull with Dirección General de Cómputo y de Tecnologías de Información y Comunicación (DGTIC-UNAM) project LANCAD-UNAM-DGTIC-290 for use of the Miztli Supercomputer.

## Figure legends

**Supplemental Figure 1.**
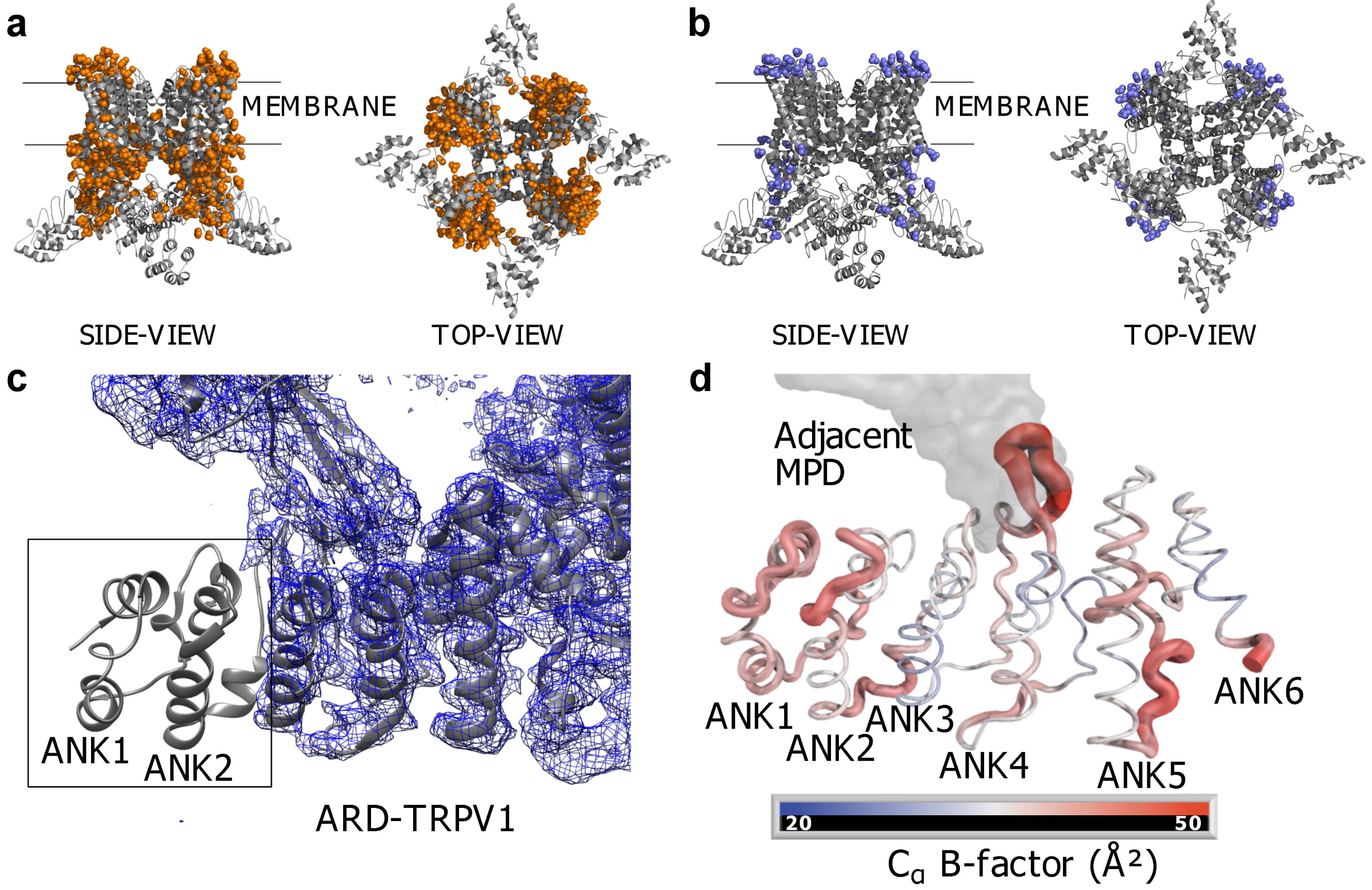
Quality of TRPV1 cryo-EM structures and crystallographic structure of ARD. (a) Side view of the missing occupancies of side chain atoms in the capsaicin-bound (CB) structure, PDBID: 2J5R. Orange spheres indicate the amino acids where the electronic density is not seen and the heavy atoms are not described; a representation by spheres shows the amino acids missing in the structure. Bottom view of the CB structure showing missing atoms in the cryo-EM structure. (b) Missing occupancies in the APO structure (3J5P), the missing occupancies vs. the CB structure are represented; the APO structure is a reliable model for coarse-grained MD. (c) Ankyrin motifs 1 (ANK1) and 2 (ANK2) do not show an electronic density in the 2J5P structure, this implies a low stability or high movement even at cryogenic temperatures. The electron density is shown in blue, while the structure of the ARD determined by X-ray diffraction is fitted to the density and show in ribbon representation. (d) Beta-factor representation in putty cartoon of the crystallographic structure of the ARD in solution (PDBID: 2PNN), code color shows red regions have poor density (assumed higher movement) and blue regions have a good density (lower movement).

**Supplemental Figure 2.**
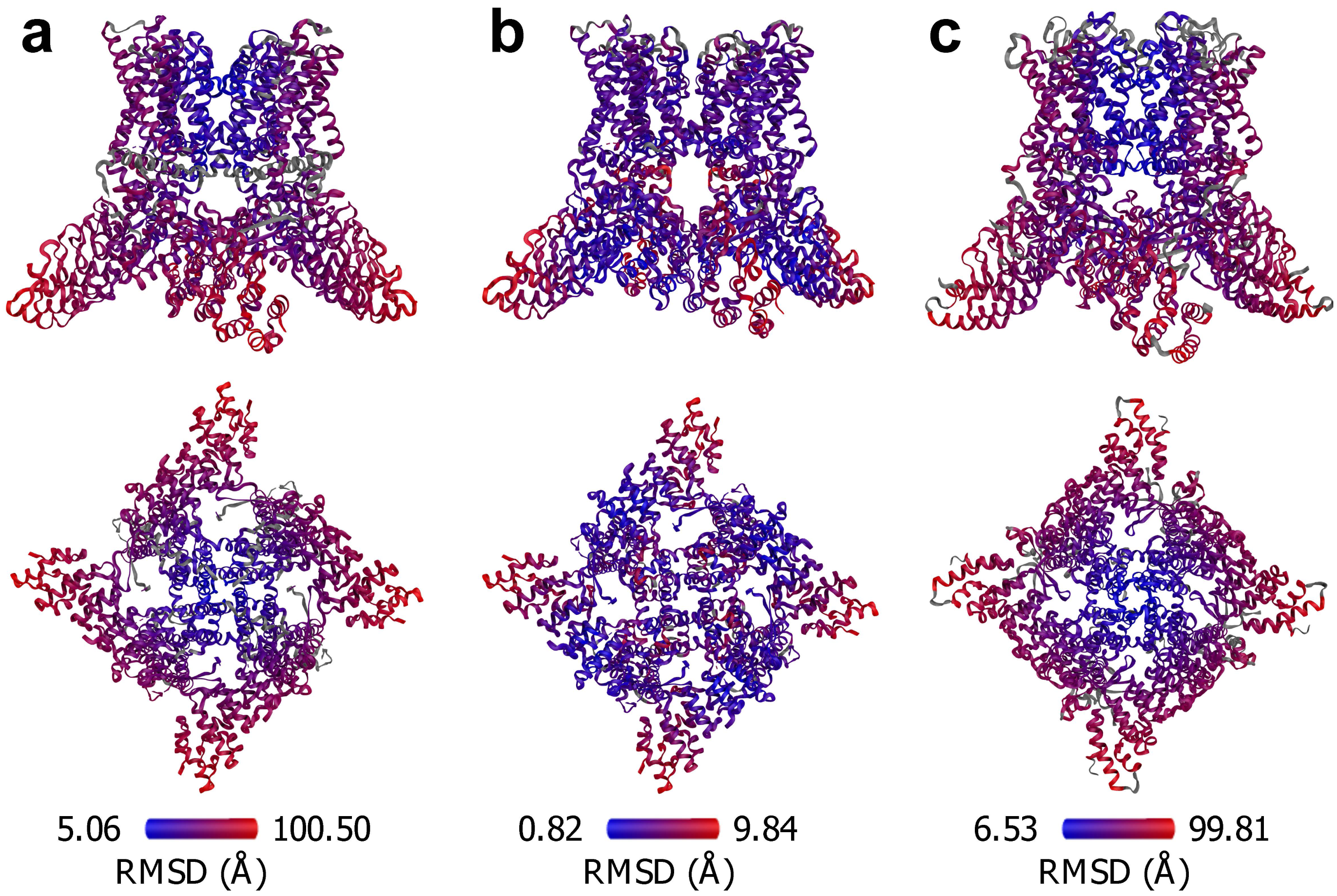
Composite alignment of closed and open structures. (a) Root mean square deviation of CB structure vs. closed structure of TRPV1 (3J5R/3J5P), the color code represents low to high RMSD by the blue to red gradient plotted in Pymol. The gray areas are not aligned by the algorithm. The values are represented over the CB structure; higher variations are seen in the cytosolic regions and the ARD presents higher movement in ANK1 and ANK2. (b) Flexible alignment of DkTx and RTX bound structure vs. closed structure (3J5Q/3J5P). The color code is as in (a). The RMSD is represented over the DkTx+TRX -bound structure. Higher values are seen in ANK 1 and ANK2 and the inner region of the TRP-box. (c) Alignment of the 2APB-bound structure of TRPV3 vs. closed structure of TRPV3 (6DVY/6DVW) represented as in (a). The main changes are similar to CB vs. closed alignment in TRPV1 (a). The color bars indicate the RMSD value scale.

**Supplemental Figure 3.**
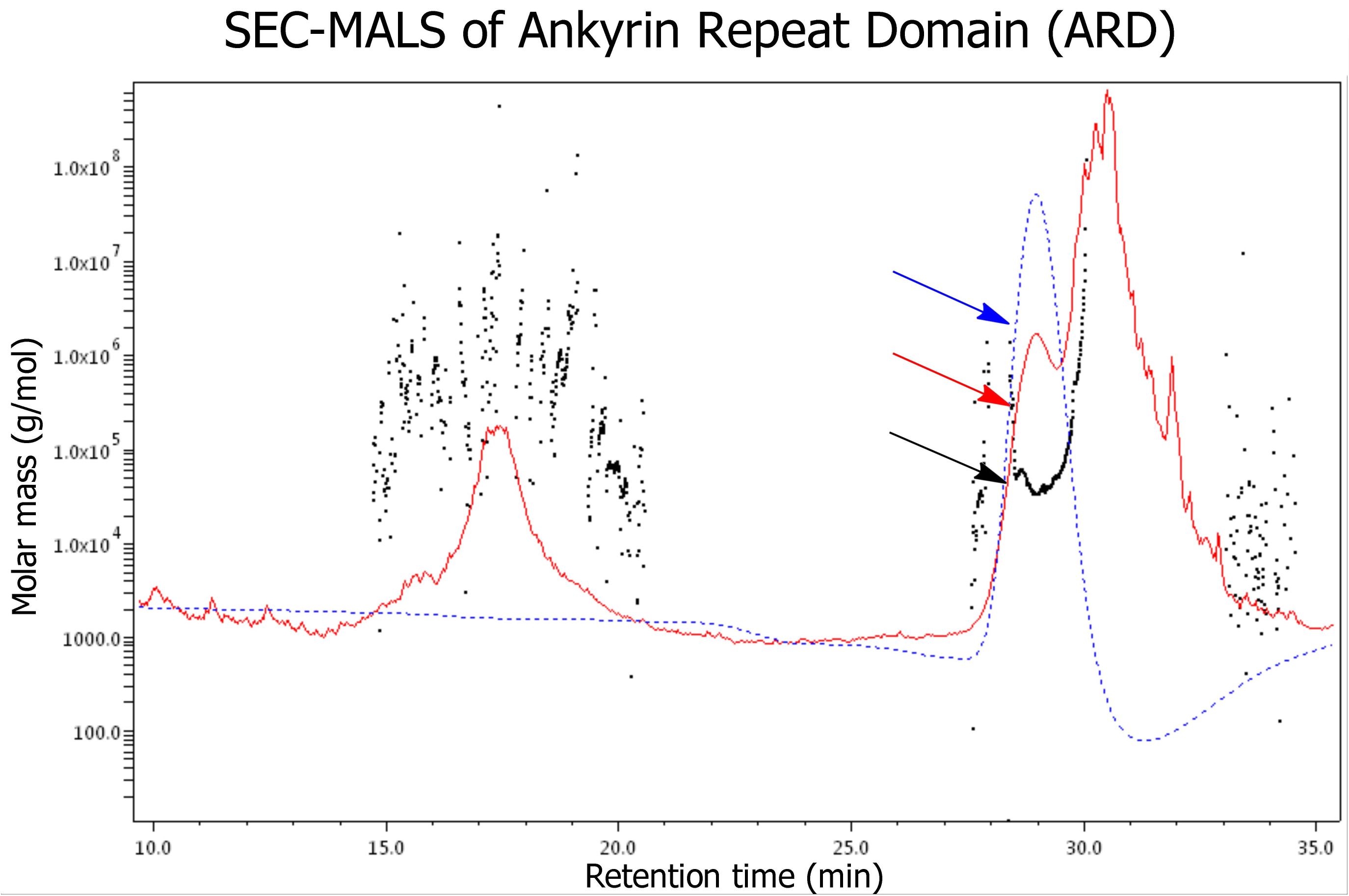
SEC-MALS of isolated ankyrin repeat domain. Chromatogram obtained in 20 mM glycine, pH 9.5, 200 mM NaCl, 25 °C. The peak (arrows) corresponds to a population with MW = 32.4 kDa and Rg = 3.1 nm. Colors correspond to light scattering (red), refractive index (blue) and molar mass (black).

## References

1. Aneiros, E., L. Cao, M. Papakosta, E.B. Stevens, S. Phillips, and C. Grimm. 2011. The biophysical and molecular basis of TRPV1 proton gating. EMBO J. 30: 994–1002.

2. Caterina, M.J., M.A. Schumacher, M. Tominaga, T.A. Rosen, J.D. Levine, and D. Julius. 1997. The capsaicin receptor: A heat-activated ion channel in the pain pathway. Nature. 389: 816–824.

3. Salazar, H., I. Llorente, A. Jara-Oseguera, R. Garcia-Villegas, M. Munari, S.E. Gordon, L.D. Islas, and T. Rosenbaum. 2008. A single N-terminal cysteine in TRPV1 determines activation by pungent compounds from onion and garlic. Nat. Neurosci. 11: 255–261.

4. Pingle, S.C., J. a. Matta, and G.P. Ahern. 2007. Capsaicin receptor: TRPV1 a promiscuous TRP channel. Handb. Exp. Pharmacol. 179: 155–171.

5. Yarmolinsky, D.A., Y. Peng, L.A. Pogorzala, M. Rutlin, M.A. Hoon, and C.S. Zuker. 2016. Coding and Plasticity in the Mammalian Thermosensory System. Neuron. 92: 1079–1092.

6. Caterina, M.J., A. Leffler, A.B. Malmberg, W.J. Martin, J. Trafton, K.R. Petersen-Zeitz, M. Koltzenburg, A.I. Basbaum, and D. Julius. 2000. Impaired nociception and pain sensation in mice lacking the capsaicin receptor. Science. 288: 306–13.

7. Huang, S.M., X. Li, Y. Yu, J. Wang, and M.J. Caterina. 2011. TRPV3 and TRPV4 ion channels are not major contributors to mouse heat sensation. Mol. Pain. 7: 37.

8. Park, U., N. Vastani, Y. Guan, S.N. Raja, M. Koltzenburg, and M.J. Caterina. 2011. TRP vanilloid 2 knock-out mice are susceptible to perinatal lethality but display normal thermal and mechanical nociception. J. Neurosci. 31: 11425–36.

9. Iida, T., I. Shimizu, M.L. Nealen, A. Campbell, and M. Caterina. 2005. Attenuated fever response in mice lacking TRPV1. Neurosci. Lett. 378: 28–33.

10. Nieto-Posadas, A., G. Picazo-Juárez, I. Llorente, A. Jara-Oseguera, S. Morales-Lázaro, D. Escalante-Alcalde, L.D. Islas, and T. Rosenbaum. 2011. Lysophosphatidic acid directly activates TRPV1 through a C-terminal binding site. Nat. Chem. Biol. 8: 78–85.

11. Ufret-Vincenty, C.A., R.M. Klein, M.D. Collins, M.G. Rosasco, G.Q. Martinez, and S.E. Gordon. 2015. Mechanism for phosphoinositide selectivity and activation of TRPV1 ion channels. J. Gen. Physiol. 145: 431–42.

12. Liao, M., E. Cao, D. Julius, and Y. Cheng. 2013. Structure of the TRPV1 ion channel determined by electron cryo-microscopy. Nature. 504: 107–12.

13. Cao, E., M. Liao, Y. Cheng, and D. Julius. 2013. TRPV1 structures in distinct conformations reveal activation mechanisms. Nature. 504: 113–8.

14. Gao, Y., E. Cao, D. Julius, and Y. Cheng. 2016. TRPV1 structures in nanodiscs reveal mechanisms of ligand and lipid action. Nature. 534: 347–51.

15. Zubcevic, L., M.A. Herzik, B.C. Chung, Z. Liu, G.C. Lander, and S.Y. Lee. 2016. Cryo-electron microscopy structure of the TRPV2 ion channel. Nat. Struct. Mol. Biol. 23: 180–186.

16. Zubcevic, L., M.A. Herzik, M. Wu, W.F. Borschel, M. Hirschi, A.S. Song, G.C. Lander, and S.-Y. Lee. 2018. Conformational ensemble of the human TRPV3 ion channel. Nat. Commun. 9: 4773.

17. Deng, Z., N. Paknejad, G. Maksaev, M. Sala-Rabanal, C.G. Nichols, R.K. Hite, and P. Yuan. 2018. Cryo-EM and X-ray structures of TRPV4 reveal insight into ion permeation and gating mechanisms. Nat. Struct. Mol. Biol. 25: 252–260.

18. Hughes, T.E.T., R.A. Pumroy, A.T. Yazici, M.A. Kasimova, E.C. Fluck, K.W. Huynh, A. Samanta, S.K. Molugu, Z.H. Zhou, V. Carnevale, T. Rohacs, and V.Y. Moiseenkova-Bell. 2018. Structural insights on TRPV5 gating by endogenous modulators. Nat. Commun. 9: 4198.

19. McGoldrick, L.L., A.K. Singh, K. Saotome, M. V. Yelshanskaya, E.C. Twomey, R.A. Grassucci, and A.I. Sobolevsky. 2018. Opening of the human epithelial calcium channel TRPV6. Nature. 553: 233–237.

20. De-la-Rosa, V., G.E. Rangel-Yescas, E. Ladrón-de-Guevara, T. Rosenbaum, and L.D. Islas. 2013. Coarse architecture of the transient receptor potential vanilloid 1 (TRPV1) ion channel determined by fluorescence resonance energy transfer. J. Biol. Chem. 288: 29506–17.

21. Rosenbaum, T., A. Gordon-Shaag, M. Munari, and S.E. Gordon. 2004. Ca2+/calmodulin modulates TRPV1 activation by capsaicin. J. Gen. Physiol. 123: 53–62.

22. Yao, J., B. Liu, and F. Qin. 2011. Modular thermal sensors in temperature-gated transient receptor potential (TRP) channels. Proc. Natl. Acad. Sci. U. S. A. 108: 11109–14.

23. Singh, A.K., L.L. McGoldrick, and A.I. Sobolevsky. 2018. Structure and gating mechanism of the transient receptor potential channel TRPV3. Nat. Struct. Mol. Biol. 25: 805–813.

24. Kasimova, M.A., A. Yazici, Y. Yudin, D. Granata, M.L. Klein, T. Rohacs, and V. Carnevale. 2018. Ion Channel Sensing: Are Fluctuations the Crux of the Matter? J. Phys. Chem. Lett. 9: 1260–1264.

25. Chuang, H., and S. Lin. 2009. Oxidative challenges sensitize the capsaicin receptor by covalent cysteine modification. Proc. Natl. Acad. Sci. U. S. A. 106: 20097–102.

26. Waterhouse, A., M. Bertoni, S. Bienert, G. Studer, G. Tauriello, R. Gumienny, F.T. Heer, T.A.P. de Beer, C. Rempfer, L. Bordoli, R. Lepore, and T. Schwede. 2018. SWISS-MODEL: homology modelling of protein structures and complexes. Nucleic Acids Res. 46: W296–W303.

27. Bordoli, L., and T. Schwede. 2012. Automated protein structure modeling with SWISS-MODEL Workspace and the Protein Model Portal. Methods Mol. Biol. 857: 107–36.

28. de Jong, D.H., G. Singh, W.F.D. Bennett, C. Arnarez, T.A. Wassenaar, L. V. Schäfer, X. Periole, D.P. Tieleman, and S.J. Marrink. 2013. Improved Parameters for the Martini Coarse-Grained Protein Force Field. J. Chem. Theory Comput. 9: 687–97.

29. Hsu, P.-C., B.M.H. Bruininks, D. Jefferies, P. Cesar Telles de Souza, J. Lee, D.S. Patel, S.J. Marrink, Y. Qi, S. Khalid, and W. Im. 2017. CHARMM-GUI Martini Maker for modeling and simulation of complex bacterial membranes with lipopolysaccharides. J. Comput. Chem. 38: 2354–2363.

30. Pronk, S., S. Páll, R. Schulz, P. Larsson, P. Bjelkmar, R. Apostolov, M.R. Shirts, J.C. Smith, P.M. Kasson, D. van der Spoel, B. Hess, and E. Lindahl. 2013. GROMACS 4.5: a high-throughput and highly parallel open source molecular simulation toolkit. Bioinformatics. 29: 845–54.

31. Van Der Spoel, D., E. Lindahl, B. Hess, G. Groenhof, A.E. Mark, and H.J.C. Berendsen. 2005. GROMACS: fast, flexible, and free. J. Comput. Chem. 26: 1701–18.

32. Sippl, M.J., and M. Wiederstein. 2012. Detection of spatial correlations in protein structures and molecular complexes. Structure. 20: 718–28.

33. Grant, B.J., A.P.C. Rodrigues, K.M. ElSawy, J.A. McCammon, and L.S.D. Caves. 2006. Bio3d: An R package for the comparative analysis of protein structures. Bioinformatics. 22: 2695–2696.

34. Gasteiger, E., C. Hoogland, A. Gattiker, S. Duvaud, M.R. Wilkins, R.D. Appel, and A. Bairoch. 2005. Protein Identification and Analysis Tools on the ExPASy Server. In: The Proteomics Protocols Handbook. Totowa, NJ: Humana Press. pp. 571–607.

35. Hellwig, N., N. Albrecht, C. Harteneck, G. Schultz, and M. Schaefer. 2005. Homo- and heteromeric assembly of TRPV channel subunits. J. Cell Sci. 118: 917–28.

36. Hinsen, K. 1998. Analysis of domain motions by approximate normal mode calculations. Proteins. 33: 417–29.

37. Skjaerven, L., S.M. Hollup, and N. Reuter. 2009. Normal mode analysis for proteins. J. Mol. Struct. THEOCHEM. 898: 42–48.

38. Inada, H., E. Procko, M. Sotomayor, and R. Gaudet. 2012. Structural and biochemical consequences of disease-causing mutations in the ankyrin repeat domain of the human TRPV4 channel. Biochemistry. 51: 6195–6206.

39. Ruan, K., and G. Weber. 1989. Hysteresis and conformational drift of pressure-dissociated glyceraldehydephosphate dehydrogenase. Biochemistry. 28: 2144–53.

40. Ferreiro, D.U., C.F. Cervantes, S.M.E. Truhlar, S.S. Cho, P.G. Wolynes, and E.A. Komives. 2007. Stabilizing IkappaBalpha by “consensus” design. J. Mol. Biol. 365: 1201–16.

41. Barrick, D., D.U. Ferreiro, and E.A. Komives. 2008. Folding landscapes of ankyrin repeat proteins: experiments meet theory. Curr. Opin. Struct. Biol. 18: 27–34.

42. Mosavi, L.K., T.J. Cammett, D.C. Desrosiers, and Z.-Y. Peng. 2004. The ankyrin repeat as molecular architecture for protein recognition. Protein Sci. 13: 1435–1448.

43. Chowdhury, S., B.W. Jarecki, and B. Chanda. 2014. A molecular framework for temperature-dependent gating of ion channels. Cell. 158: 1148–1158.

44. Zhang, F., A. Jara-Oseguera, T.-H. Chang, C. Bae, S.M. Hanson, and K.J. Swartz. 2018. Heat activation is intrinsic to the pore domain of TRPV1. Proc. Natl. Acad. Sci. U. S. A. 115: E317–E324.

45. Hryc, C.F., D.-H. Chen, P. V. Afonine, J. Jakana, Z. Wang, C. Haase-Pettingell, W. Jiang, P.D. Adams, J.A. King, M.F. Schmid, and W. Chiu. 2017. Accurate model annotation of a near-atomic resolution cryo-EM map. Proc. Natl. Acad. Sci. U. S. A. 114: 3103–3108.

46. Lishko, P. V., E. Procko, X. Jin, C.B. Phelps, and R. Gaudet. 2007. The Ankyrin Repeats of TRPV1 Bind Multiple Ligands and Modulate Channel Sensitivity. Neuron. 54: 905–918.

47. Dhaka, A., V. Uzzell, A.E. Dubin, J. Mathur, M. Petrus, M. Bandell, and A. Patapoutian. 2009. TRPV1 is activated by both acidic and basic pH. J. Neurosci. 29: 153–8.

48. Parra, R.G., R. Espada, N. Verstraete, and D.U. Ferreiro. 2015. Structural and Energetic Characterization of the Ankyrin Repeat Protein Family. PLoS Comput. Biol. 11: e1004659.

49. Sanchez-Moreno, A., E. Guevara-Hernandez, R. Contreras-Cervera, G. Rangel-Yescas, E. Ladron-de-Guevara, T. Rosenbaum, and L.D. Islas. 2018. Irreversible temperature gating in trpv1 sheds light on channel activation. Elife.: 251124.

50. Schumacher, M.A., I. Moff, S.P. Sudanagunta, and J.D. Levine. 2000. Molecular cloning of an N-terminal splice variant of the capsaicin receptor. Loss of N-terminal domain suggests functional divergence among capsaicin receptor subtypes. J. Biol. Chem. 275: 2756–62.

